# SVFX: a machine-learning framework to quantify the pathogenicity of structural variants

**DOI:** 10.1101/739474

**Authors:** Sushant Kumar, Arif Harmanci, Jagath Vytheeswaran, Mark B. Gerstein

## Abstract

A rapid decline in sequencing cost has made large-scale genome sequencing studies feasible. One of the fundamental goals of these studies is to catalog all pathogenic variants. Numerous methods and tools have been developed to interpret point mutations and small insertions and deletions. However, there is a lack of approaches for identifying pathogenic genomic structural variations (SVs). That said, SVs are known to play a crucial role in many diseases by altering the sequence and three-dimensional structure of the genome. Previous studies have suggested a complex interplay of genomic and epigenomic features in the emergence and distribution of SVs. However, the exact mechanism of pathogenesis for SVs in different diseases is not straightforward to decipher. Thus, we built an agnostic machine-learning-based workflow, called SVFX, to assign a “pathogenicity score” to somatic and germline SVs in various diseases. In particular, we generated somatic and germline training models, which included genomic, epigenomic, and conservation-based features for SV call sets in diseased and healthy individuals. We then applied SVFX to SVs in six different cancer cohorts and a cardiovascular disease (CVD) cohort. Overall, SVFX achieved high accuracy in identifying pathogenic SVs. Moreover, we found that predicted pathogenic SVs in cancer cohorts were enriched among known cancer genes and many cancer-related pathways (including Wnt signaling, Ras signaling, DNA repair, and ubiquitin-mediated proteolysis). Finally, we note that SVFX is flexible and can be easily extended to identify pathogenic SVs in additional disease cohorts.

## Introduction

Large-scale whole-genome sequencing is providing high-resolution maps of genomic variation in various disease-specific studies^1–4^. These studies have created extensive catalogs of genomic alterations that comprise single nucleotide changes (SNVs or SNPs), insertions and deletions (INDELs, ranging between 1-50 bp), and structural variations (SVs, which exceed 50 bp). SVs are often classified as imbalanced or balanced based on their effect on the copy number profile. Imbalanced SVs result in copy number changes through large deletions, duplications or insertions. In contrast, balanced SVs (such as translocations and inversions) do not alter the copy number profile of an individual. Despite their relatively lower frequencies, SVs contribute more nucleotide-level changes than aggregated frequencies of SNVs/SNPs and INDELs^5^.

As a result of their large size, SVs play a vital role in the progression of various diseases, including cancer, intellectual disabilities, and neurodegenerative diseases^4^. In the context of cancer, these rearrangements often lead to the removal or fusion of genes and their cis-regulatory elements, thereby disrupting essential functions, including cell growth, differentiation, signaling, and apoptosis^6^. Despite their important roles in various diseases, ascertaining the pathogenicity and establishing mechanistic links between SVs and disease progression remains challenging^7^. This challenge is exacerbated by difficulties associated with accurate identification of SVs and their precise breakpoint^8^.

Prior studies aimed at quantifying the pathogenicity and ascertaining the roles of genomic variations in disease have primarily been limited to point mutations and small INDELs^9–13^. In contrast, only a handful of studies have aimed to evaluate the molecular consequence of SVs^14^. Initial attempts to characterize the molecular impact of SVs were limited to annotating genes that overlap with germline SVs, without assigning pathogenicity scores. A recent study^14^ has leveraged genome-wide per-base pathogenicity scores^9^ (initially designed for measuring the impact of single nucleotide changes) to assign impact scores for germline SVs. Despite these early efforts, there is a clear need for a systematic framework to understand the molecular and functional consequences of SVs and their roles in human disease.

To address this challenge, we present an integrative supervised machine learning framework (SVFX) to assign pathogenic scores to somatic and germline SVs. We hypothesized that the underlying genomic and epigenomic features of pathogenic SVs are very different from those of benign SVs. Moreover, these differences can be sensitively detected only in a tissue-specific context. Thus, we built machine learning models that assign a pathogenic score by comparing genomic and tissue-specific epigenomic features of a given SV to known benign SVs. Our framework is highly flexible and can be applied to identify pathogenic somatic and germline SVs in cancer as well as other diseases. Toward this end, we utilized high-quality somatic and germline SVs from the Pan-Cancer Analysis of the Whole Genomes (PCAWG) Project^15^, Genome Sequencing Program (GSP) and the 1000 Genomes (1KG) Project^5,16^ to train our machine learning model. Additionally, we employed tissue-specific epigenomic data from Epigenome Roadmap^17^, various genomic element annotations^18,19^, and cross-species conservation metrics^20^ to build our machine learning models.

Overall, our approach achieved high accuracy in discriminating pathogenic somatic SVs from benign variants for large deletions (mean AUC of 0.865) and duplications (mean AUC 0.835) across multiple cancer types. Additionally, our germline models attained good accuracy in identifying pathogenic germline SVs in cancer as well as cardiovascular disease. In particular, our somatic model identified pathogenic deletions and duplications in cancer genome that are enriched among key pathways and biological processes, including cell cycle regulation, cell differentiation, and signal transduction. Additionally, for somatic models where we excluded conservation and known cancer gene annotation as features, we found that high-impact (pathogenic) SVs tend to influence highly conserved regions of the genome and are enriched among known cancer genes. This observation provided further evidence for the robustness of our approach in identifying pathogenic SVs. Finally, we annotate and discuss examples of somatic SVs that were identified as highly pathogenic using our method.

## Results

### Training dataset and SV impact workflow

For each disease cohort, we built separate somatic and germline models. In somatic SV models, a training set consisted of cancer and control (i.e., benign SVs from the 1KG project) SVs (**Fig 1a**). For the germline model, we sub-sampled germline SVs for each cancer cohort such that the number of germline SVs in the disease set is same as the common SVs (global allele frequency > 0.5%) in the 1KG SV dataset^5^. Additionally, the CVD cohort in our study had a unique advantage of being a careful case-control study. Thus, instead of using common 1KG SVs as a benign variant, we utilized common SVs belonging to the control group from this study as benign SV dataset.

**Fig 1.**
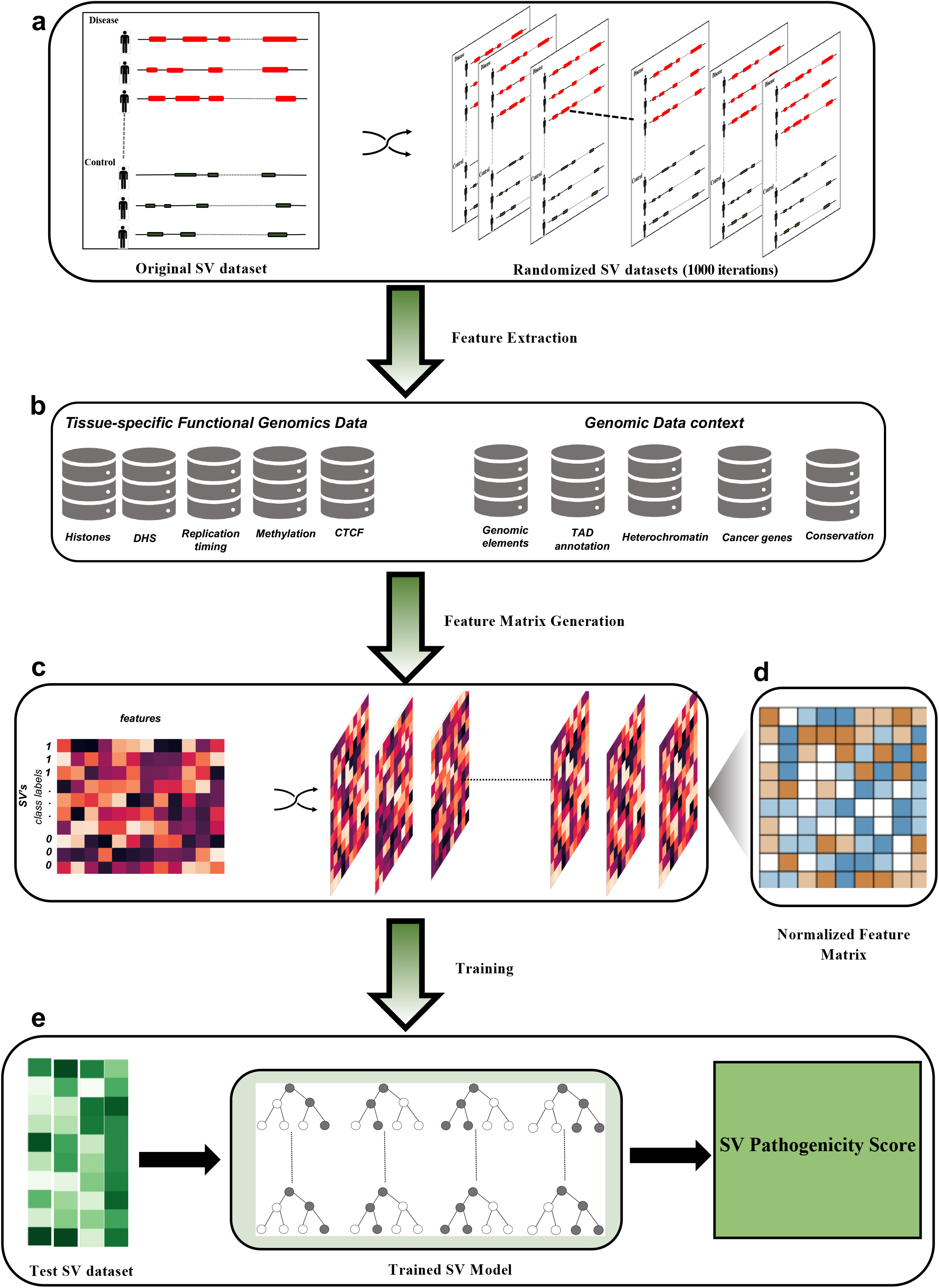
Machine learning based workflow of SVFX to identify pathogenic structural variations: a) The original SV dataset consist of disease/case and control SVs. In our somatic model disease SVs correspond to somatic SVs found in a cancer cohort and control SVs correspond to SVs found in the one thousand genome project (1KG SVs). We randomly select SVs from 1KG SV dataset such that the number of somatic and control SV matches. Similarly, for our germline model we have disease germline SVs identified in specific disease cohort and control SVs, which correspond to common SVs in 1KG SV dataset. For both germline and somatic models, we generate thousand random iteration of original disease and control dataset. These permuted SVs are later utilized for generating Z-score normalized feature matrix.

Previous works^21–24^ have shown that distribution of somatic and germline SVs depends on the complex interplay between different mechanistic biases that originates from the underlying chromosome conformations^23^, DNA accessibility^22,25^, functional annotation^5^, methylation profile^24^ and cross-species conservation. For instance, disease SVs disrupts topologically associated domains which influence the gene-enhancer interaction leading to various diseases^23^. Similarly, methylation status^24^ and DNA accessibility^25^ of the genomic regions have been previously associated with the emergence of somatic and germline structural variations. Furthermore, disruption of coding and non-coding genomic elements (including promoters, UTRs, and enhancers) along with highly conserved genomic regions are likely to play a critical role in disease progression^5^. Despite these strong correlations, for many diseases the exact mechanism of how a pathogenic SV drives disease progression remains elusive. Thus, in this work we adopted a data driven approach where we built agnostic machine learning models incorporating various genomic and epigenomic features underlying SVs. Further, we hypothesized that the genomic and epigenomic profile of pathogenic SVs are very distinct from benign ones.

Accordingly, we built feature matrices for our somatic and germline models, where each row corresponded to an SV and each column to a distinct feature. These feature matrices consist of important epigenomic features, including average histone mark signals, methylation levels, CTCF signals, open chromatin marks, and replication timing data that overlapped with SVs in the disease and the benign datasets. Furthermore, we also integrated relevant genomic element annotations, including the fraction of overlap between SVs and the coding region, 3’ and 5’ UTR, splice sites and promoter region of genes in the human genome. The feature matrices also captured additional annotations including TAD boundaries definition, heterochromatin regions, fragile sites, sensitive sites, and ultra-conserved regions in the genome (**Fig 1b**).

A unique challenge in the feature-based representation of SVs is that in disease cohorts (especially somatic ones) and 1KG dataset, they exhibit an apparent disparity in their length distributions. These differences in length distributions for disease and benign SVs are likely to influence various feature values implicitly. Thus, to avoid any bias in training features, we uniformly shuffled SVs in disease cohorts as well as for the benign SV dataset to generate null distributions of feature values (**Fig 1c**). We next transformed each feature value in the original feature matrices (consisting of disease and benign SVs) to obtain Z-score normalized feature matrices using the null distribution of each corresponding element (**Fig 1d**). While Z-score based normalization compensates for the length bias, we would like to make sure that our models assign high pathogenic scores to extremely long SVs such as chromosomal arm-level variants. Thus, we appended the length of each SVs as an explicit feature in our Z-score normalized feature matrices.

For each disease cohort, we utilized these updated Z-score normalized feature matrices for training supervised machine learning models using random forest algorithm for somatic and germline SVs separately (**Fig 1e**). Finally, we validated these trained models using ten-fold cross-validation, as well as in the independent test dataset for each cohort.

### Accuracy assessment of somatic cancer models

We applied our method to quantify the pathogenic score of somatic SVs in six different cancer cohorts. These cohorts included breast, ovarian, liver, esophageal, stomach, and skin cancers. We selected these cohorts based on available tissue-specific epigenomic data, and also because these cancer types consist of a significant number of SVs (which are needed for training and testing the model). Subsequently, we evaluated whether these models could identify pathogenic somatic SVs from benign ones. Intuitively, one would expect that our somatic model should assign high impact scores to cancer SVs, whereas low pathogenicity score would be assigned to benign 1KG SVs. Moreover, we would expect that SVs with high pathogenic scores would play the role of cancer drives, whereas low score SVs were likely to be passenger variants with little or no consequence on tumor progression. We quantitatively assessed this using 10-fold cross-validation, as well as among independent test dataset for each cancer cohort. Briefly, we measured the average areas under the receiver operator characteristic (auROC) and the precision-recall (auPR) curves.

Overall, our somatic models for both deletions and duplications performed very well. For somatic deletions with the 10-fold cross-validation strategy, we observed the mean auROC and auPR values across all six cancer types to be 0.861 and 0.892, respectively (**Fig 2a, supplement Fig S1a**). In contrast, the mean auROC and auPR values for somatic duplication models across six cancer cohorts were 0.835 and 0.87, respectively (**Fig 2b, supplement Fig S2a**). In addition to the 10-fold cross-validation, we also assessed the performance of our somatic deletion and duplication models in an independent test dataset. Overall, we observed comparable performance, with an average auROC value of 0.865 and 0.835 across six cancer types for deletions and duplications, respectively (**Fig 2c and Fig 2d**). For the independent test data, the average auPR values were also very similar to those using 10-fold cross-validation. Our models achieved a mean auPR value of 0.87 and 0.89 across six cancer types for deletions and duplications, respectively (**supplement Fig S1b & S2b**). Furthermore, we also quantified the pathogenic score for large deletions and duplications that are predicted to be driver events on a pan-cancer level using a recurrence-based analysis^26^. As expected, our workflow assigned a high pathogenic score (average score greater than 0.9 across different somatic models) to each putative driver event.

**Fig 2.**
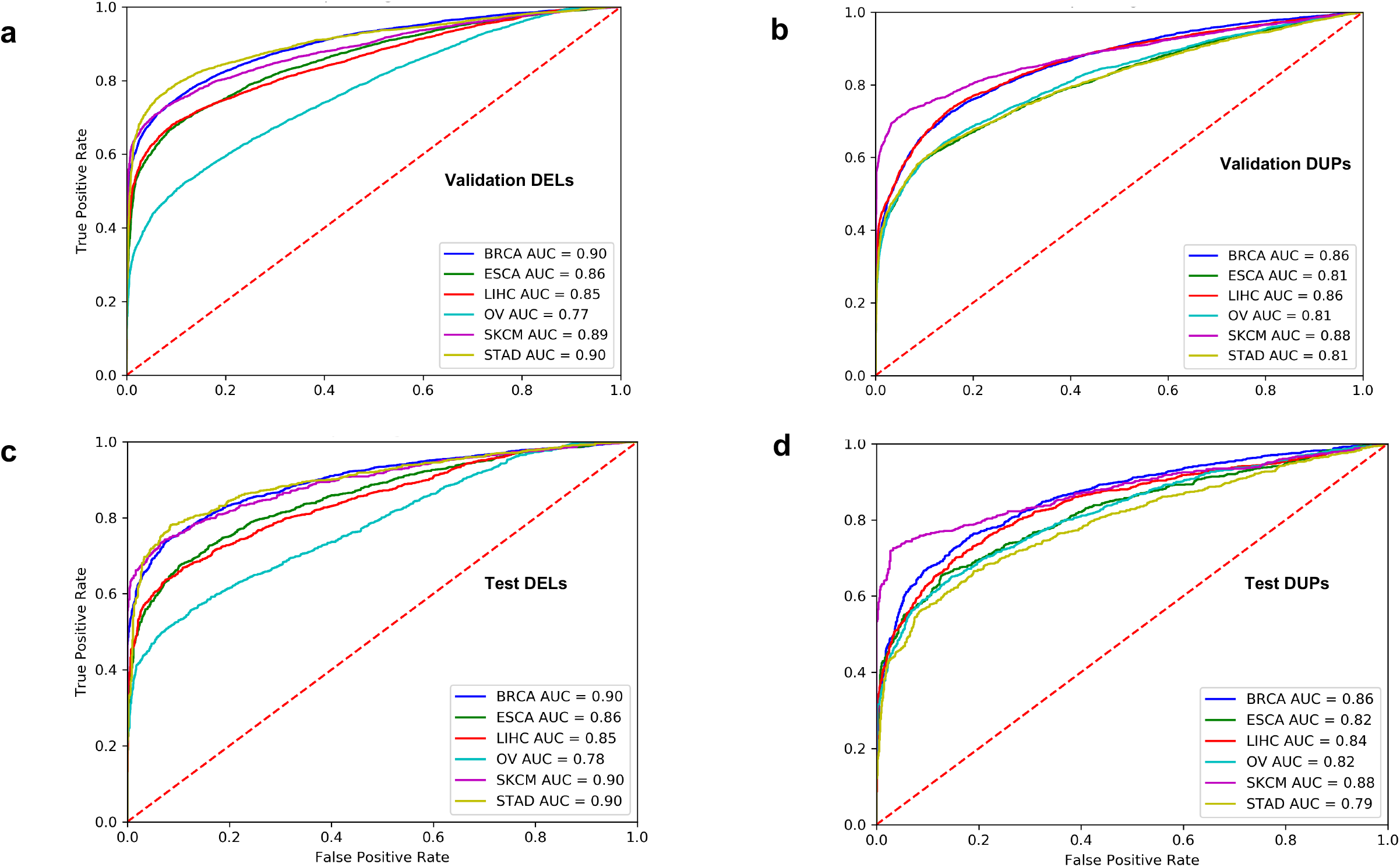
Performance evaluation for somatic models to predict pathogenic SVs in various cancer types: This figure presents Area Under the receiver Operating Curve(auROC) based on the validation datasets for large deletions (panel a) and duplications (panel b) in six different cancer cohorts including breast adenocarcinoma (BRCA), esophageal carcinoma (ESCA), liver (LIHC), ovary (OV), skin melanoma (SKCM) and stomach (STAD) cancers. Similarly, auROC plots for test datasets associated with large deletions (panel c) and duplications (panel d) in six different cancer cohorts.

Finally, we also evaluated the contribution of each feature to the performance of our somatic deletion and duplication models. We observed that SV length and overlap with ultra-conserved regions are the most significant contributors to the predictive performance of the model (**supplement Fig S7**).

Additionally, predictability of somatic deletion models depends on other noncoding and epigenomic features, including overlap with 3’ and 5’ UTR, sensitive regions and H3K4me3 signals suggesting an essential influence of SVs on cis-regulatory elements. Similarly, the predictive performance of somatic duplication models primarily depends on overlap with known cancer genes, heterochromatin annotation, UTRs, and sensitive regions (**supplement Fig S8**).

### Accuracy assessment of germline models

In addition to somatic models, we built germline SV models to identify pathogenic germline SVs in six cancer cohorts. We assessed the predictive accuracy of our germline models in cancer cohorts using 10-fold cross-validation and independent test dataset. Similar to somatic deletions, we observed good performance for our germline deletion models in cancer cohorts. Using 10-fold cross-validation, the mean auROC and auPRC values across six cancer types were 0.79 and 0.74, respectively (**supplement Fig S3a & S4a**). Additionally, we observed very similar auROC and auPR values among different cancer types. Among our test datasets, the average auROC and auPR values across different cancer cohorts were approximately 0.8 and 0.75, respectively (**supplement Fig S3b & S4b**). We note that the ovarian cancer cohort primarily drove the small improvement of the model in the test dataset with auROC and auPR values of 0.86 and 0.84, respectively (**supplement Fig S3b & S4b**).

We note that our current framework is highly flexible and can be easily applied to identify pathogenic germline SVs in other diseases. For example, we used our framework to build a germline model for cardiovascular disease cohort belonging to the BioMe biobank program (included in the GSP project^27^). Overall, our germline deletion model in this cardiovascular disease (CVD) cohort achieves good predictive accuracy with mean auROC and auPR values of 0.77 and 0.74, respectively (**supplement Fig S5a & S6a**). Similarly, our model performs very well to identify pathogenic deletions in the testing dataset for this cohort with mean auROC and auPR values of 0.76 and 0.84, respectively (**supplement Fig S5b & S6b**).

Similar to somatic models, we also quantified the prediction contribution for each feature in our germline deletions and duplications model. We observed that the length of SVs provides a maximum contribution to our cancer germline predictions. Additionally, we found a substantial contribution from sensitive regions, cancer gene overlap, H3K9me3, and 3’ UTR overlap (**supplement Fig S9**). In contrast, the overlap of SVs with TAD boundary annotations had the most significant contribution for assigning pathogenicity score for the CVD cohort (**supplement Fig S10**). This observation is consistent with previous studies highlighting the role of germline SVs in various diseases through disruption of the 3D-genome structure.

### Somatic model evaluation and gene enrichment analyses

We performed various analyses to investigate the biological validity and robustness of our approach for quantifying the pathogenicity of cancer SVs. For instance, we carried out a separate round of investigations in which we excluded cross-species conservation scores and overlap fraction with ultraconserved and sensitive regions from our models. We computed the pathogenic score of each SV using these modified models and correlated them with the average PhyloP score for genomic regions overlapping with the SV. Relative to low pathogenic score SVs, somatic SVs with higher pathogenic scores should intuitively be expected to overlap with more conserved regions of the genome. Indeed, we observed that highly pathogenic (SV pathogenic score >= 0.9) deletions and duplications exhibited higher conservation scores compared to benign deletions and duplications (SV pathogenic core <= 0.2). This observation was highly significant for both deletions and duplications (**Fig 3a**).

**Fig 3.**
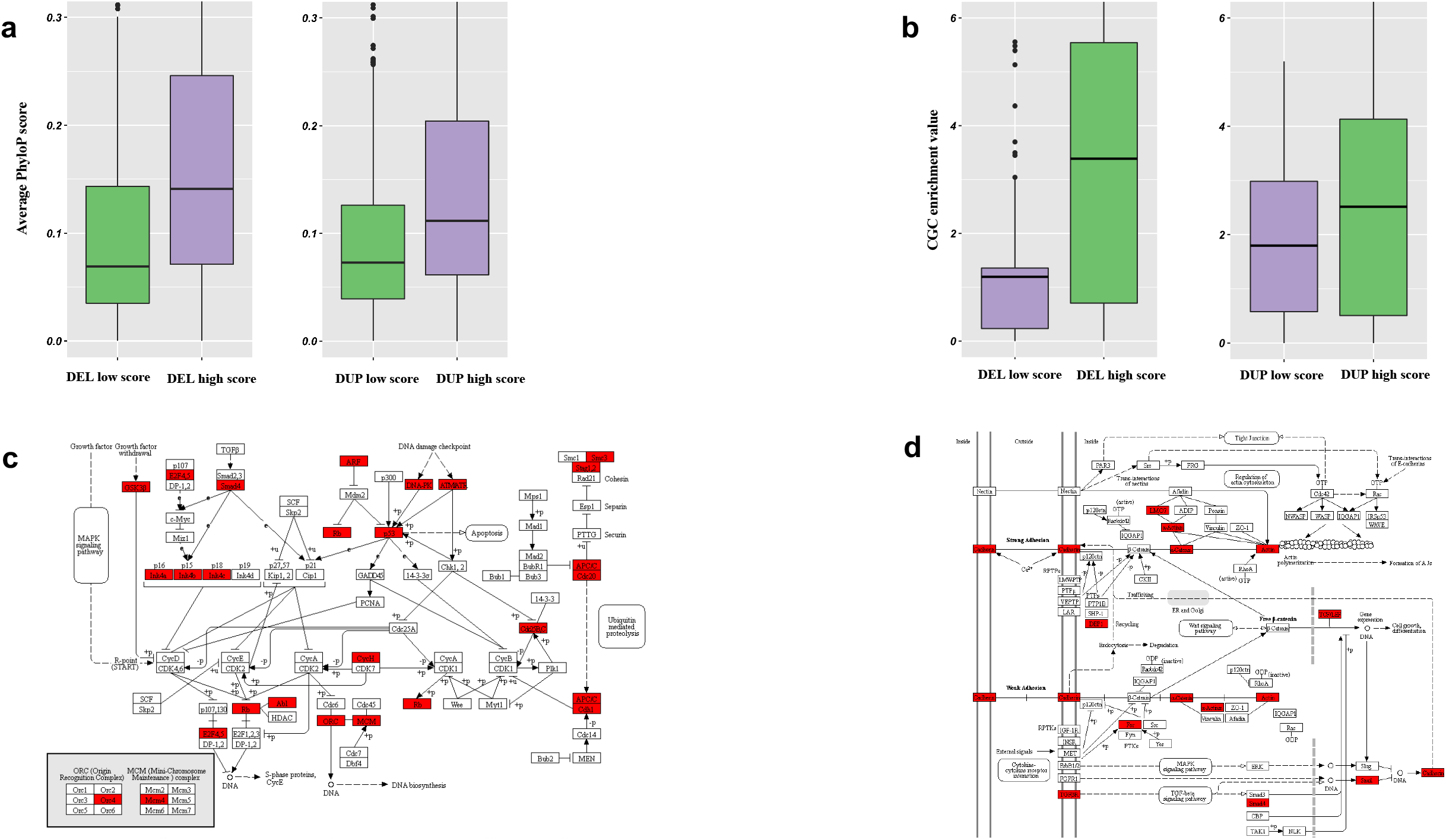
Orthogonal biological validations of somatic models in cancer: a) This plot presents mean conservation score comparison for genomic regions that overlap with predicted highly pathogenic deletions against benign deletions (left panel) and duplications (right panel) for a model where conservation was excluded from the original model. b) This plot presents cancer gene enrichment value for coding regions that overlap with predicted highly pathogenic deletions against benign deletions (left panel) and duplications (right panel) for a model where overlap fraction with cancer genes was excluded from the original model. c) Example of the ubiquitin-mediated proteolysis pathway that is enriched among genes that are affected by highly pathogenic deletions on the pan-cancer level. Genes that are influenced by highly pathogenic deletions in this pathway are highlighted in red. d) Example of the adherens junction pathway which is enriched among genes that are affected by highly pathogenic duplications on the pan-cancer level. Genes that are influenced by highly pathogenic duplications in this pathway are highlighted in red.

**Fig 4.**
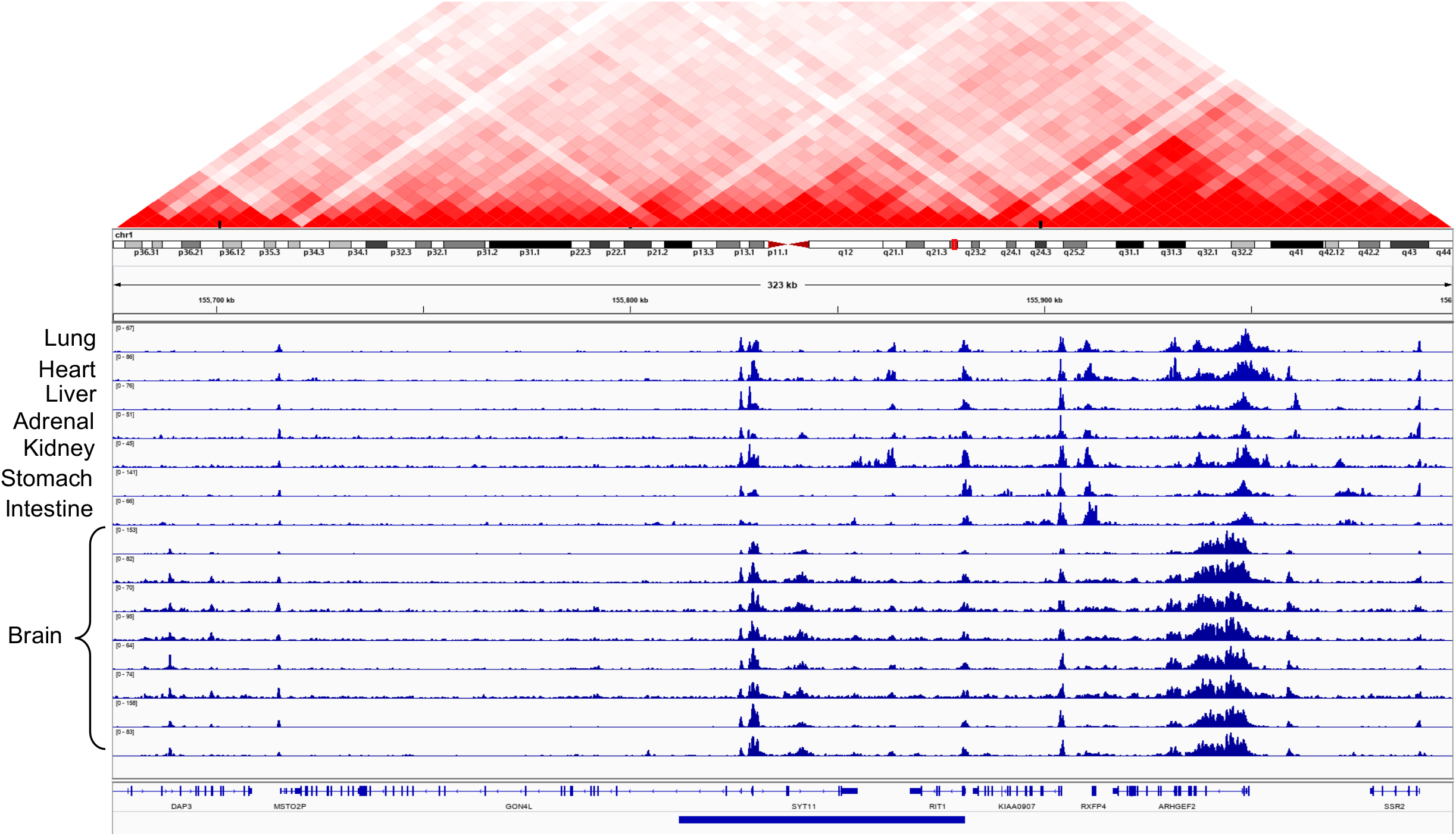
Example of highly pathogenic cancer deletion that influence coding and regulatory elements in the genome: In this figure we present highly pathogenic deletion that disrupts entirely or partially the coding and regulatory regions of three distinct genes including *RIT1, SYT11, and GON4L.* The regulatory elements are marked by peaks observed in the histone mark (H3K27ac) signals across multiple tissues. We also plot the HIC matrix to show TAD boundaries disrupted by this deletion.

Next, we assessed whether our machine learning approach assigns a high pathogenic score to SVs that are enriched among known cancer genes. As with the analysis detailed above, we excluded the known cancer gene overlap as a feature to generate modified random forest models for this analysis. We recomputed the pathogenic score for each SV using these modified models, and then classified SVs as high- and low-pathogenic SVs based on the thresholds detailed above. Subsequently, we quantified the enrichment of known cancer genes in high- and low-pathogenic SV groups. As expected, we observed more substantial enrichment of known cancer genes among highly pathogenic deletions and duplications compared to those with lower pathogenic scores (**Fig 3b**). As with our conservation analysis, differences in enrichment between these SV groups were found to be highly significant.

Moreover, we used our original model to identify highly pathogenic deletions and duplications (SV pathogenic score >= 0.9) on the pan-cancer level. We then identified all coding genes that entirely or partially overlapped with these pathogenic deletions and duplications. Using this overlapping gene list, we performed ontology and pathway enrichment analysis. We observed that pathogenic deletions influence those genes that are enriched for vital biological processes, including signal transduction, cell cycle, post-translational modification, and DNA repair. Pathway-level analyses indicate that these pathogenic deletions affect critical pathways that involve Wnt and Ras signaling, cellular senescence, transcriptional regulation, and ubiquitin-mediated proteolysis (**supplement Fig S11 & supplement data tables 2-3**). In particular, we highlight genes (in red) that are deleted by highly pathogenic SVs and which are involved in the ubiquitin-mediated proteolysis pathway (**Fig 3c**). Our results are consistent with prior studies, which have shown that disruption of these pathways can drive tumor progression^28^.

Similarly, highly pathogenic duplication influenced genes that are enriched for cell differentiation, development, signal transduction, and various metabolic processes. Pathway-level enrichment analysis suggests that such duplicated genes play a pivotal role in Tyrosine receptor kinase signaling, post-translational protein modifications, membrane trafficking, and Wnt signaling pathway (**supplement Fig S12 & supplement data tables 4-5**). We highlight a set of genes, including Cadherin, Actin, SMAD4, DEP and TGF beta receptor that is affected by highly pathogenic duplication and plays a vital role in the adherens junction pathway (**Fig 3d**). The adherens junction pathway maintains homeostatic cell signaling, and its disruption is known to drive breast cancer progression^29^.

### Case studies highlighting high impact somatic deletion and duplication

Our machine learning framework can clearly distinguish between pathogenic cancer SVs and benign SVs. Based on the pathogenicity score, we highlight examples of somatic deletion and duplications that are predicted to be highly pathogenic in different cancer cohorts. Overall, we found that many deletions and amplifications with high pathogenic scores overlapped with regulatory regions in the genome. To visually inspect the effect of these variants, we used the H3K27ac histone modification from multiple tissues generated by the Roadmap Epigenome Mapping Consortium^17^. This particular histone modification marks the presence of cis-regulatory elements such as promoters and enhancers in the genome. Overall, we observed that these example SVs influence regulatory elements that are active in multiple tissues, as reflected in the conserved H3K27ac signal profiles. Presumably, these conserved regulatory elements play an essential role in gene regulation, and thus their disruption through deletions or duplications is likely to be highly pathogenic. Moreover, we also used the Hi-C contact matrix to inspect the chromatin structure around these deletions and amplifications^30^.

For instance, **Fig 4** shows annotation of a high impact deletions that is also recurrent across multiple cancer types. As expected, this particular deletion overlapped with several non-coding elements, completely engulfing two genes (SYT11 and RIT1), and partially overlapping with the first exon of another gene (GON4L). The RIT1 gene encodes for a protein that plays a crucial role in the RAS/MAPK pathway and regulates the cellular signals required for cell proliferation and differentiation. RIT1 gene belongs to the RAS family of an oncogene. A previous study^31^ indicates an association between RIT1 gene inactivation and lymphoma progression. Similarly, the GON4L gene is a transcription regulator that plays a vital role in cell division and differentiation. In particular, GON4L gene-based transcription regulation is essential for B cell development and differentiation^32^. Moreover, the SYT11 gene encodes a protein that facilitate calcium-signal dependent membrane trafficking. In addition to affecting coding region of these genes, this particular deletion also engulfs many cis-regulatory elements and thus influence their long-range interaction. In particular, we observe perturbation of numerous 3-dimensional interactions (shown by the Hi-C contact matrix above) that a deleted enhancer makes within the vicinity of the locus.

Finally, we describe an example of a highly pathogenic somatic duplication (**Fig 5**), which directly overlapped with the SETD3 gene, a histone methyltransferase that is implicated in many diseases, including cancers^33^. The SETD3 gene also plays a role in cell cycle regulation, dell death, and chromosomal translocation^34,35^. Furthermore, the overexpression of the SETD3 gene leads to cell proliferation and tumor growth in liver cancer cells^36^. In addition to SETD3, this particular amplification also affects the CCNK gene, which has a vital role in transcriptional regulation. The chromatin structure from Hi-C data showed interactions of the regulatory element effected by this amplification with the nearby genes that include YY1. The YY1 gene plays a dual role in activating and repressing of a large number of gene promoters. Overall, these examples highlight the efficacy of our approach in identifying highly pathogenic structural variants. Furthermore, they provide essential insights into higher-order regulatory interactions that are affected by some of these variants.

**Fig 5.**
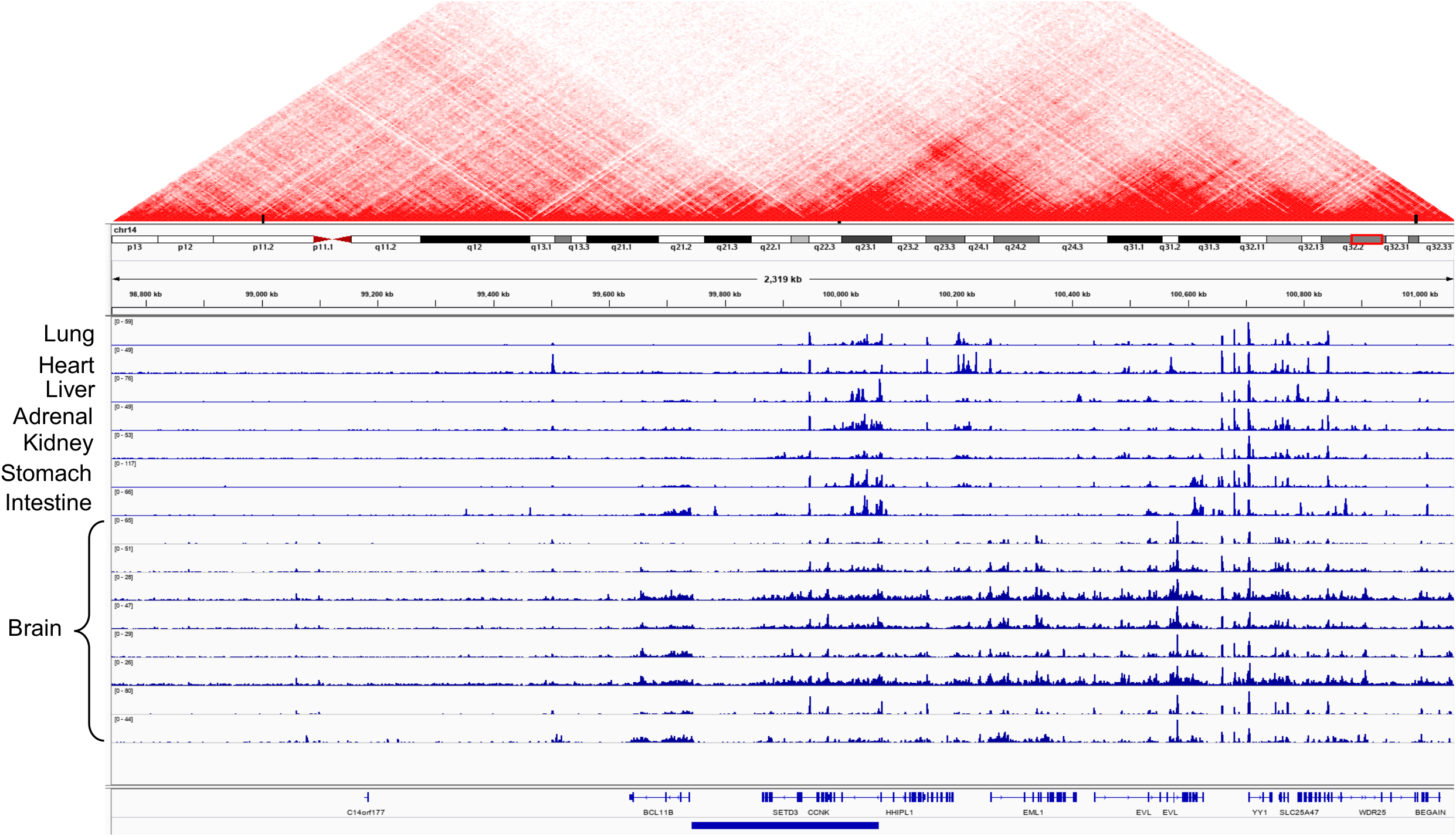
Example of highly pathogenic cancer duplication that influence coding and regulatory elements in the genome: This panel presents highly pathogenic duplication that influences coding and regulatory elements for multiple genes, including *BCL11B, SETD3, CCNK, and HHIPL1.* Similar to pathogenic deletions, this panel also displays the HIC profile to highlight TAD boundaries that are disrupted by this highly pathogenic duplication.

## Discussion

One of the fundamental goals of population-level^37^ and disease-specific sequencing studies^27,38,39^ has been to identify causal SNPs and INDELs in a large pool of candidate variants. Such efforts resulted in multiple tools and metrics to prioritize SNPs and INDELs. However, large SVs represent an essential set of variations that influence linear as well as the three-dimensional genome structure. These alterations can perturb protein-coding regions and cis-regulatory elements alike, and they are often involved in various diseases, including cancer^7^. Despite altering a considerable fraction of the genome, there have been relatively few systematic studies to prioritize and identify pathogenic SVs. A lack of such efforts can be partially attributed to challenges associated with accurate identification of SVs and their precise breakpoints using the short-read sequencing technologies^8^. However, with the development of better tools and methods for SV discovery using short- and long-read techniques^8^, we anticipate that generating a high-resolution map of genomic rearrangements will soon become routine. Thus, it is becoming essential to develop new methodologies for evaluating the pathogenicity of SVs.

In this work, we present a new machine learning-based framework to assess the pathogenicity of SVs in disease cohorts. Although the current approach utilizes cancer and cardiovascular disease data, it can easily be extended to any other disease study. Overall, our method very accurately identifies highly deleterious SVs and distinguishes them from low scoring benign SVs. For somatic deletions in cancer, we achieved a mean auROC value of 0.865 across multiple cancer types (with a maximum of 0.9), including breast, melanoma, and stomach cancers. Similarly, for somatic duplications, we achieved similar performance, with a mean auROC value of 0.835 across multiple cancer types. Likewise, the performance of our cancer germline deletion model was good (with a mean auROC of 0.8) across six cancer cohorts. Additionally, auROC values of the germline models for different cancer cohorts were remarkably similar. We expect that including additional high-quality common SVs in our training dataset will further improve the discriminative performance of the germline model. Finally, we also applied our framework to assign a pathogenic score to germline SVs in a case-control study of cardiovascular disease. In this cohort, our germline model also achieved good predictive accuracy (with a mean auROC of 0.76), which is comparable to those of somatic models.

We also built multiple versions of our original somatic models to evaluate the biological validity and robustness of our approach. For instance, we excluded the cross-species conservation score and other related annotation features from our initial model to re-compute the SV pathogenicity score of our training and test SV dataset. As expected, we observed a higher average conservation score for highly pathogenic SVs compared to low scoring benign SVs. Similarly, we built yet another version of our original model, in which we removed annotations for known cancer genes. We observed that high scoring SVs identified from these models were significantly enriched among known cancer genes compared to low scoring SVs. This observation further suggests that our machine learning framework is robust and assign biologically intuitive pathogenic scores. Finally, we performed pathway and ontology enrichment analyses of genes that overlapped with high scoring SVs, as identified with our original model. We observed an enrichment of genes in many cancer-related pathways, including Wnt signaling, Ras signaling, DNA repair, cell differentiation, and ubiquitin-mediated proteolysis. These results further support the biological validity of our approach for assigning pathogenic scores to cancer-associated SVs.

As briefly noted above, our machine learning framework is flexible and can be easily extended to assign a pathogenic score for SVs in other disease-specific whole-genome studies, including autism and neuro-developmental diseases. Additionally, we note that our current framework primarily focuses on identifying pathogenic deletions and duplications. However, it can be readily extended to detect pathogenic inversions and translocations in these diseases. In summary, despite their crucial role in various diseases, few approaches are currently available for interpreting and prioritizing SVs. At a per-nucleotide level, SVs contribute far more substantial variation in an individual genome than other mutations. However, SVs are often neglected as a consequence of the technical challenges associated with their identification and interpretation. We address this challenge by building a new framework that utilizes tissue-specific genomic and epigenomic features to quantify the pathogenicity of SVs. Identification of such pathogenic SVs along with deleterious point mutations and INDELs will facilitate a complete understanding of the biology for various diseases.

## Method

Structural variant coordinates were first gathered from the Pan-Cancer Analysis of Whole Genomes (PCAWG) project^15,26^ and the 1KG SV datasets. Then, to form a control group without a bias for length, random structural variant coordinates were generated across the genome, each equivalent in length to the variant from the cancer and 1KG source. 1KG SVs were assumed to be benign, and the variants from the cancer source, while not all deleterious, were expected to contain some set of harmful variants, which we wanted to identify through our method. We followed a similar approach for germline SVs as well. However, for cancer germline SVs, we utilized common 1KG SVs as benign variants. In contrast, for the CVD cohort, we leveraged SVs present in the control group as the benign variant set.

For each SV (identified solely by coordinates), a variety of features were calculated and compiled into a feature matrix. Three categories of features were selected: Tissue-specific functional genomics data, various annotation metrics (**Supplement data table 1**), and conservation scores. Annotation overlaps were calculated as the percentage of the variant that overlapped with any region in a given annotation dataset - for example, given a 10,000-nucleotide variant and a set of coordinates corresponding to TADs, if 5,000 of the nucleotides in the variant lied in one of the TADs, then the overlap metric would be 0.5. For tissue-specific epigenomic and functional genomics data-based features, we divided SVs into windows of 10 base pairs length and computed the features over these windows. For instance, given an SV [*a, b*] which starts at genomic position a and ends at position b, we divided the interval into 10 base pair bins, i.e., 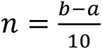 bins. For the *i^th^* bin (*n*≥ *i* ≥ 1), we computed the total signal values for each functional genomics and epigenomic dataset within the bin. Subsequently, we calculated the average of these value for each dataset over all 10bp bins that overlap with a given SV. Putting these together, the total set of features can be summarized as:

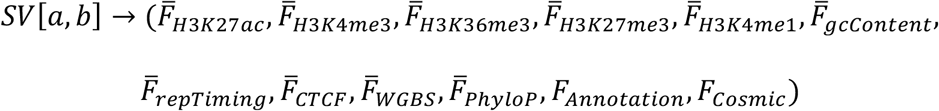

where 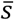 denotes the average signal over all the 10-bp bins within [*a, b*]. Overall, we computed 22 features and use them to build the model. As discusses earlier, we used these features because prior studies have shown strong correlation with subset of these features and distribution of structural variations in the genome^21–25^. Additionally, our annotation-based features are likely to capture properties of the coding and non-coding functional elements in the genome.

Subsequently, all features were normalized by Z-Score transformation, and class labels were assigned (1 for all variants from the disease dataset, and 0 for 1KG SVs). While all disease SVs are not deleterious (in fact, a minority were expected to be), the rationale behind the labeling method was that benign variants would mostly share characteristics with 1KG SVs. Thus, our model would only predict a variant’s class to be 1 with high confidence if the variant was very different in character from a benign variant. Moreover, we also appended length of SVs in Z-score transformed feature matrices for training and testing of machine learning models.

Once the feature matrix was compiled and normalized, the data was used to train ten random forest models. Each model trained on a disjoint 10% of the data. Then, each model predicted a probability for the remaining 90% of the data for a class label of 1 (in other words, that the variant was from the disease associated SV data set). Then, the nine probabilities for each variant were averaged to produce one final score, meant to reflect the probability that the variant was a member of the disease-associated dataset, and thus, through ordering variants by these scores, a ranking of variants could be constructed. Variants with very high probabilities, near the top of the ranking, had characteristics that were very different from the set of “benign” 1KG variants, while variants with low (around 0.5 and below) probabilities had features that were virtually indistinguishable from those of benign variants. The model’s hyper-parameters (maximum depth of each tree in the forest, number of trees in the forest, and minimum number of leaves required to split an internal node) were tuned to maximize the Area Under the Receiver-Operator Curve (auROC) and the Area Under the Precision Recall Curve (auPRC). The source code for the SVFX workflow is available on the project’s Github page (https://github.com/gersteinlab/SVFX).

## Downstream analyses

In order to perform orthogonal validation, we modified the original feature matrix to generate two modified models. In one such model, we removed average cross-species conservation (PhyloP) score and the overlap fraction of SVs with ultra-conserved and sensitive regions in the human genome. Similarly, in a different model, we removed features capturing overlap fraction of SVs with known cancer genes as defined in the cancer gene census database. For both these modified models, we followed the same procedure of Z-score based feature normalization, training and testing. For the model without conservation score, we defined highly pathogenic SVs based on pathogenicity score threshold above 0.9 and benign SVs with pathogenicity score below 0.2. For pathogenic and benign classes of SVs, we then computed the average conservation score by taking the mean value of nucleotide-level PhyloP score for regions overlapping with a given SV.

Similarly, for the model without cancer gene annotation, we applied the same SV impact thresholds to classify SVs into benign and pathogenic groups. For each group of SVs, we computed the fraction of overlap between known cancer genes and member SVs for different cancer types. For enrichment calculation, members of pathogenic and benign SV group were permuted thousand times across the genome. For each cancer gene, we computed the fraction of nucleotide overlapping with original and permuted SVs to calculate a Z-score based enrichment score. Subsequently, we compared these Z-score enrichment scores to measure differences between pathogenic and benign SVs. Finally, we also calculated the gene ontologies and pathway enrichments of genes that partially or completely overlapped with highly pathogenic SVs. Pathway enrichment was done for KEGG as well as the reactome database.

## Acknowledgements

We acknowledge support from the NIH and the AL Williams Professorship funds. We are thankful to the members of the PCAWG SV working group for generating the variant calls. We are also grateful to the Center for Common Disease, and the Genome Sequencing Program consortium members for creating SV calls for the CVD cohort used in this study.

