## Supplemental Figures for "SVFX: a machine-learning framework to quantify the pathogenicity of structural variants"

### Supplementary Figures

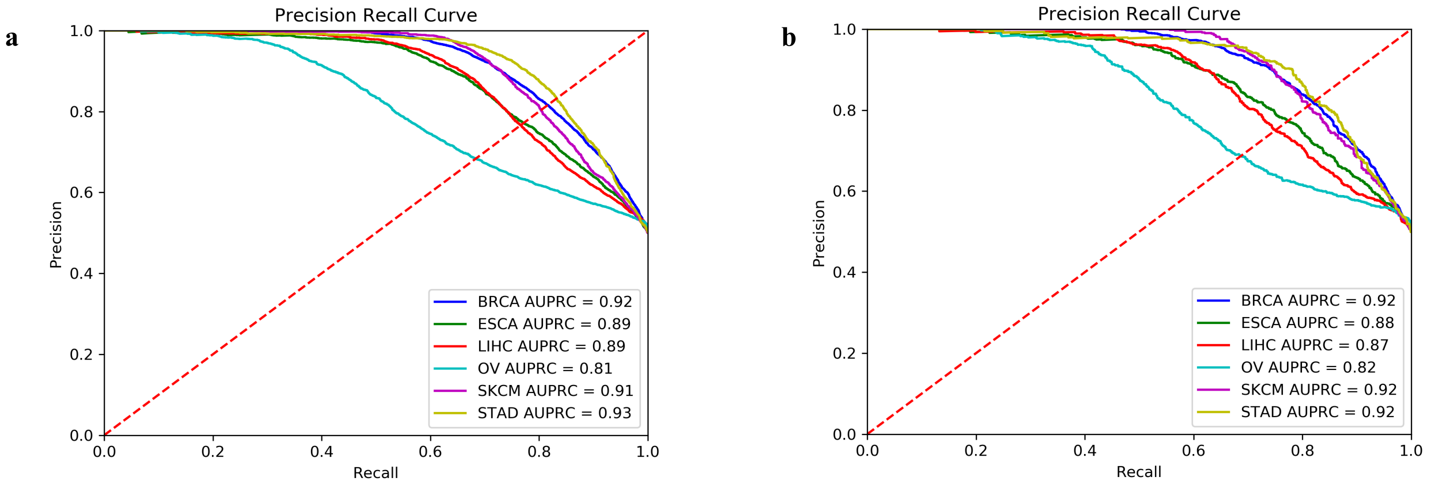


**SI FIG1:** *Area under Precision Recall Curves(auPRC) for large somatic deletions belonging to six cancer cohorts using the validation and testing datasets*. a) auPRC plots for somatic DELs in six cancer cohorts using 10-fold cross validation approach, b) auPRC plot for somatic deletions in six distinct cohorts using independent testing datasets that were not used to train random forest models.

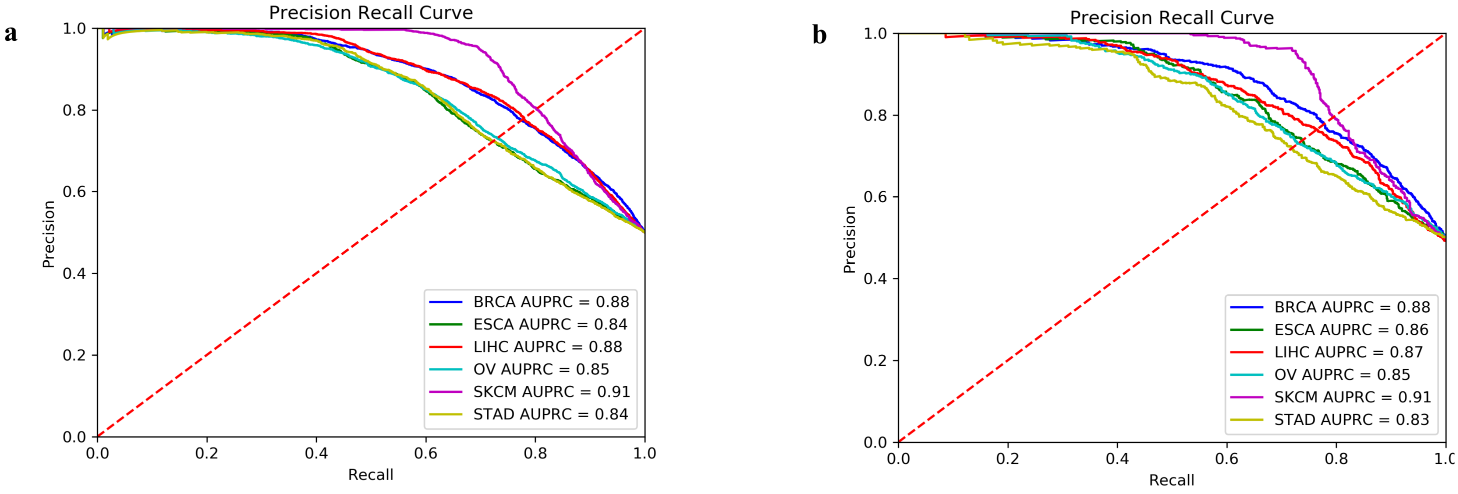


**SI FIG2:** *Area under Precision Recall Curves(auPRC) for somatic duplications belonging to six cancer cohorts using the validation and testing datasets.* a) auPRC plots for somatic duplications in six cancer cohorts using 10-fold cross validation approach, b) auPRC plot for somatic duplications in six distinct cohorts using independent testing datasets.

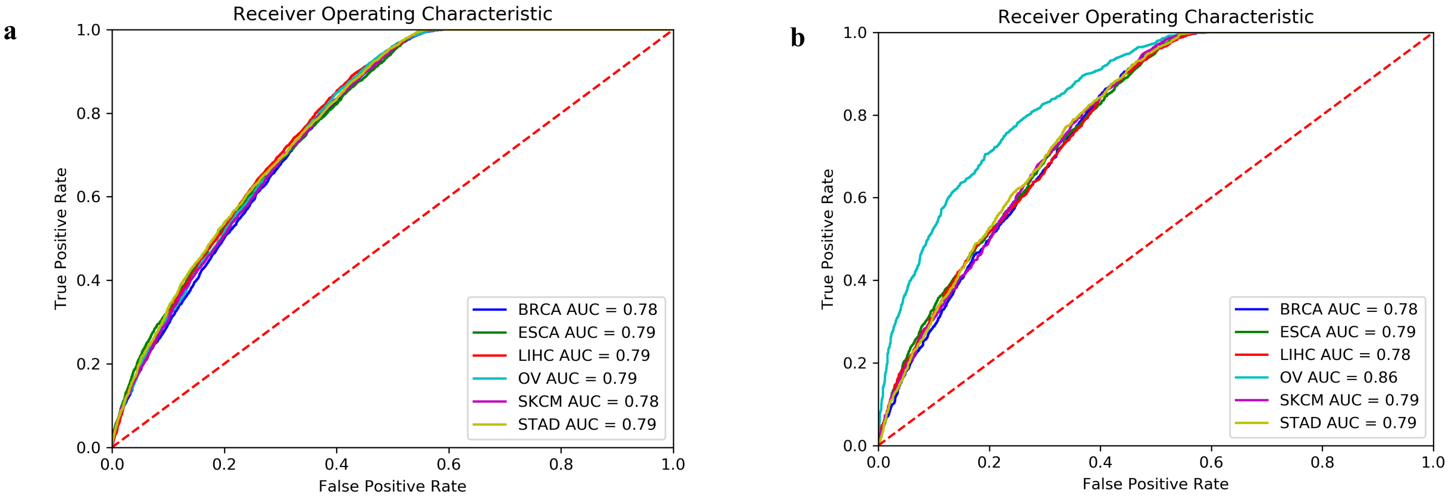


**SI FIG3:** *Area under Receiver Operating Characteristics (auROC) for germline deletions belonging to six cancer cohorts using the validation and testing datasets.* a) auROC plots for germline deletions in six cancer cohorts using 10-fold cross validation approach, b) auROC plot for germline deletions in six distinct cohorts using independent testing datasets that were not used to train random forest models.

a

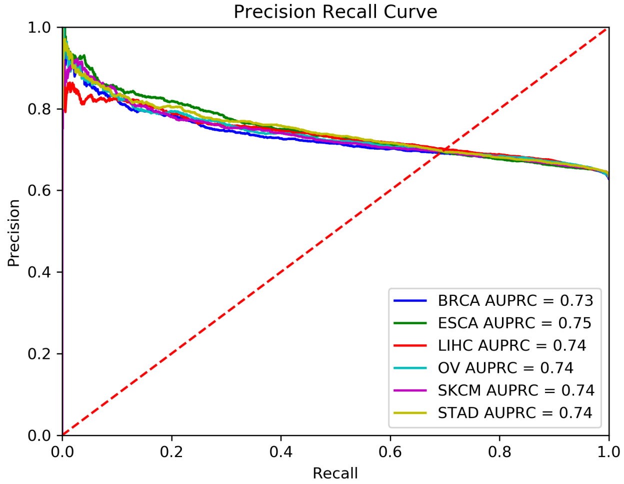


b

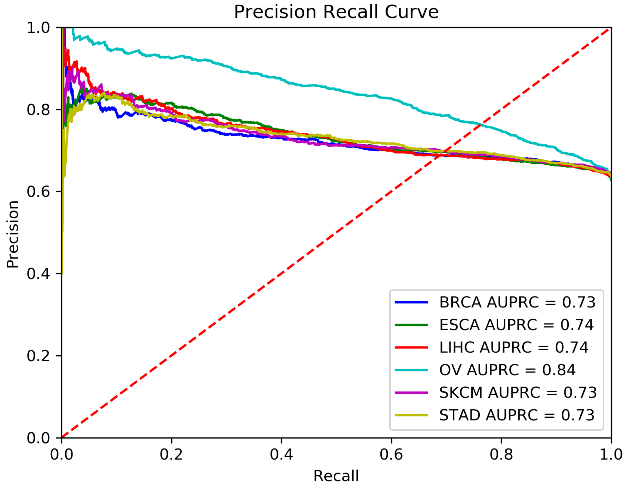


**SI FIG4:** *Area under Precision Recall Curves(auPRC) for germline deletions belonging to six cancer cohorts using the validation and testing datasets.* a) auPRC plots for germline deletions in six cancer cohorts using 10-fold cross validation approach, b) auPRC plot for germline deletions in six distinct cohorts using independent testing datasets.

a

b

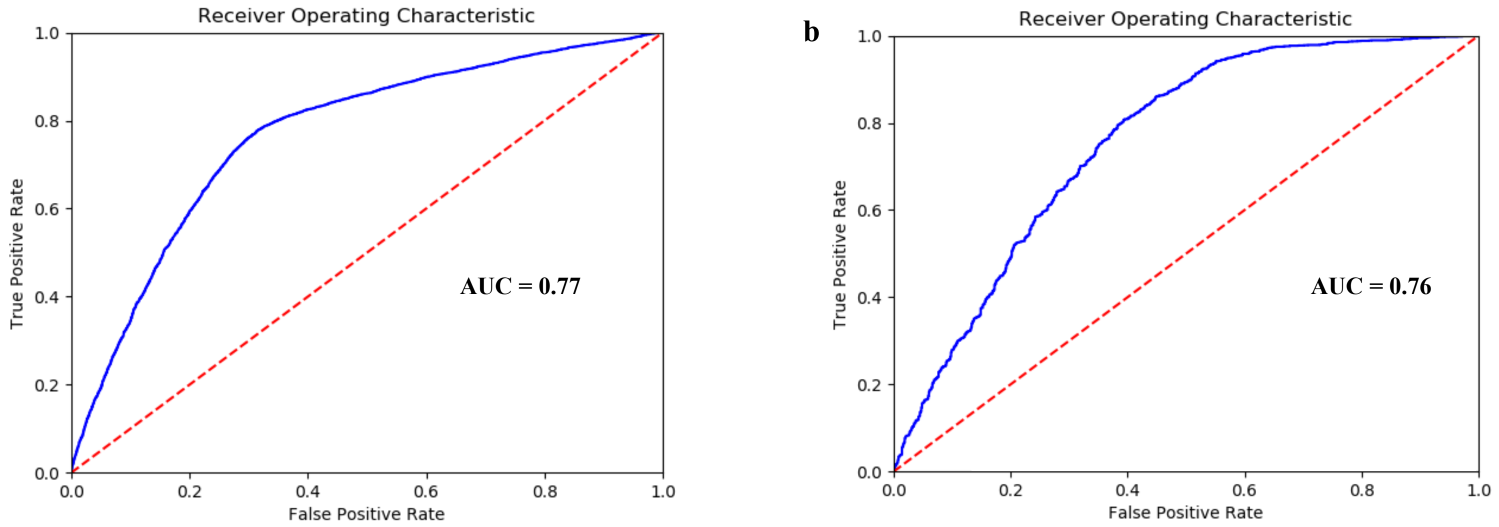


**SI FIG5:** *Area under Receiver Operating Characteristics (auROC) for germline deletions belonging to cardiovascular cohort using the validation and testing datasets.* a) auROC plots for germline deletions in the cardiovascular cohort based on 10-fold cross validation approach, b) auROC plot for germline deletions in the cardiovascular cohort using independent testing datasets.

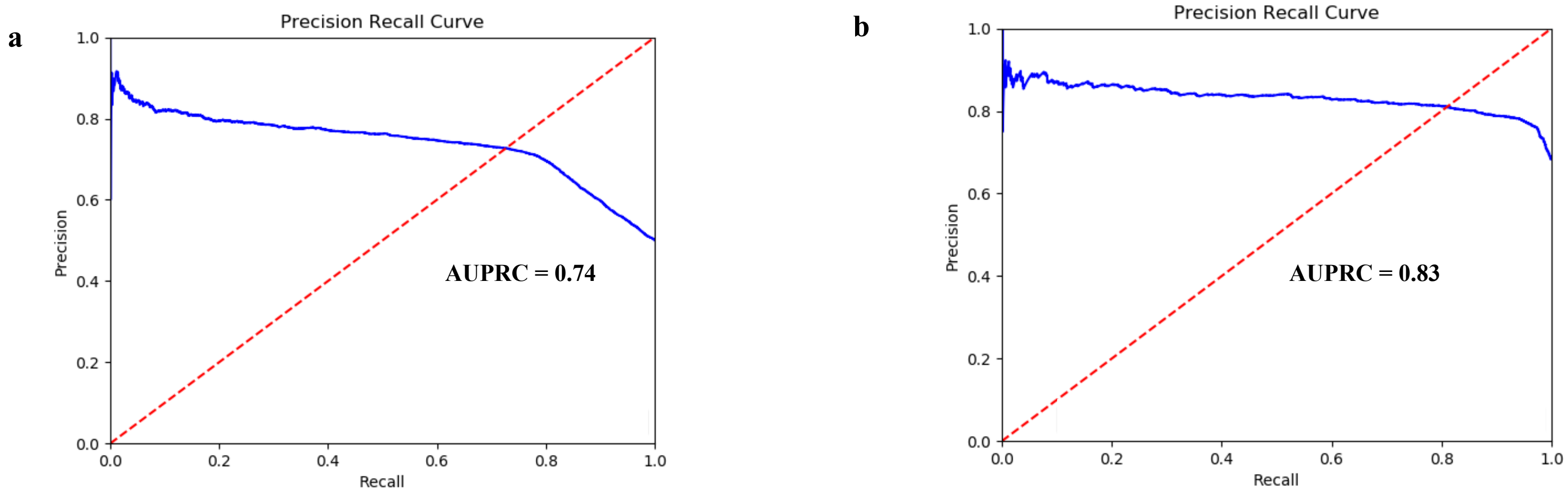


**SI FIG6:** *Area under Precision Recall Curves(auPRC) for germline deletions in the cardiovascular disease cohort using the validation and testing datasets.* a) auPRC plots for germline deletions in the cardiovascular disease cohort using the 10-fold cross validation approach, b) auPRC plot for germline deletions in the cardiovascular disease cohort using independent testing datasets.

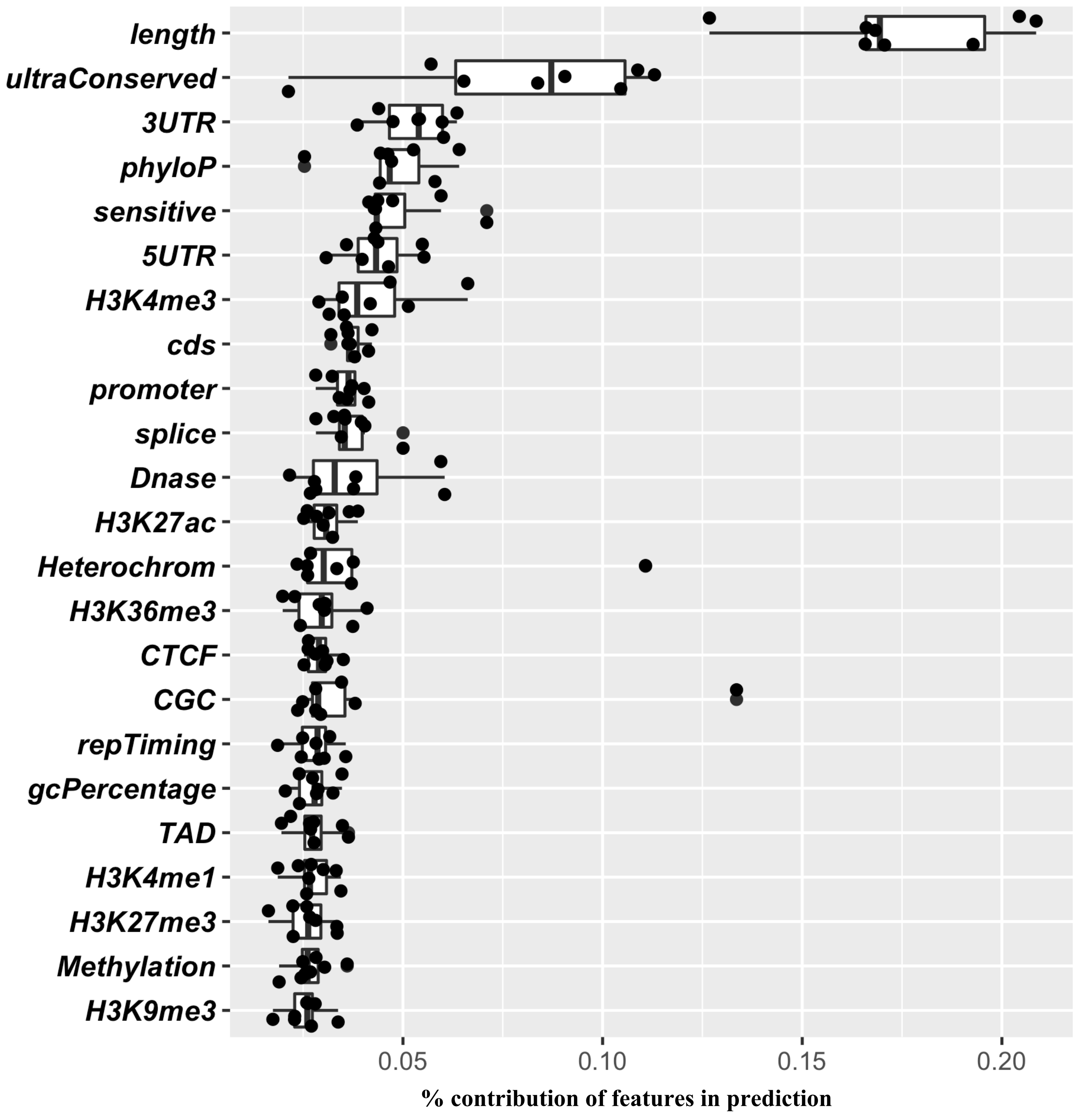


**SI FIG7:** This Figure presents the percentage contribution for distinct features in the overall predictability of somatic deletion models for six cancer cohorts. Each data point in the boxplot corresponds percentage contribution of a feature in the overall performance in a cancer cohort.

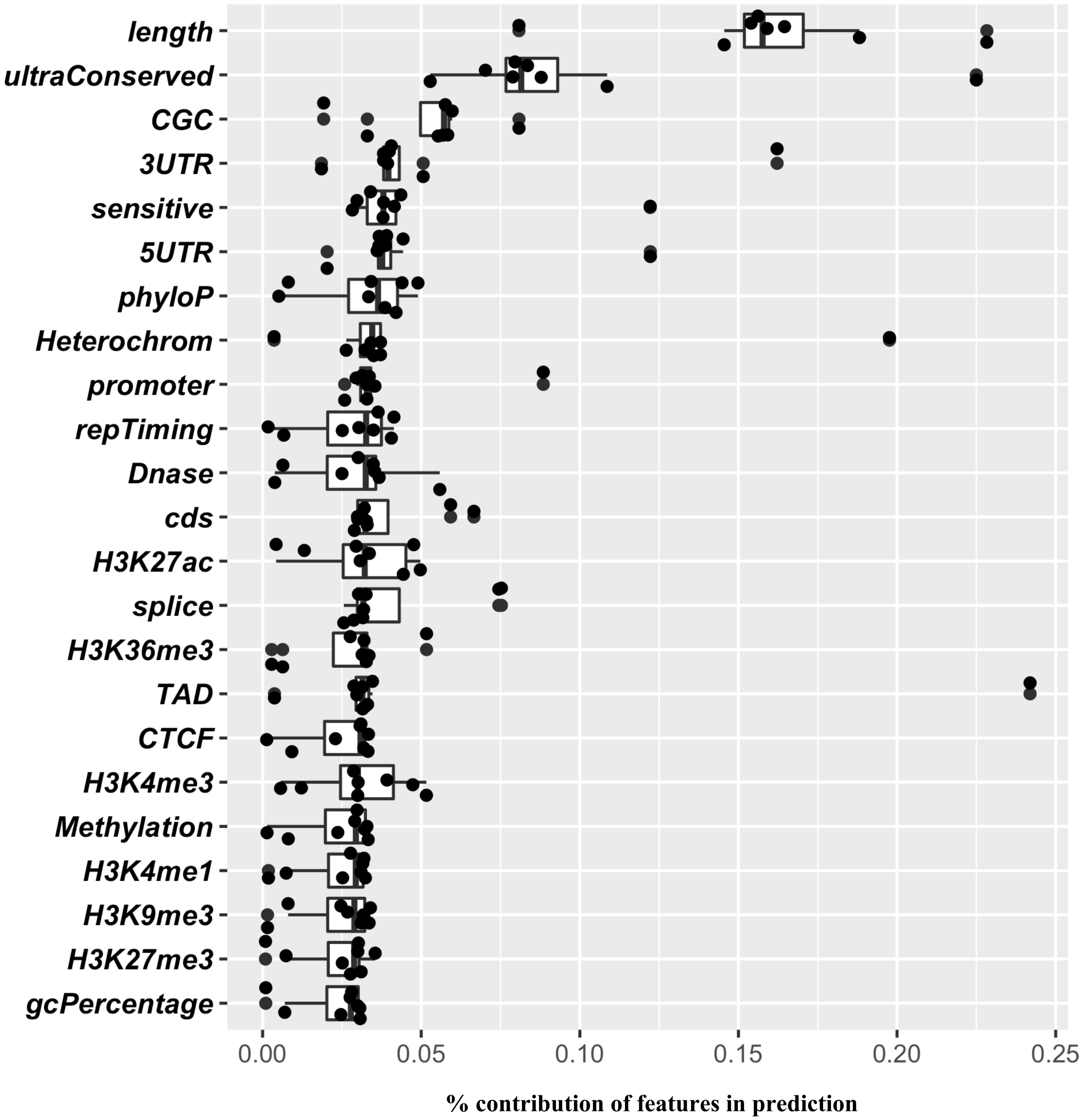


**SI FIG8:** This Figure presents the percentage contribution for distinct features in the overall predictability of somatic duplication models for six cancer cohorts. Each data point in the boxplot corresponds percentage contribution of a feature in the overall performance in a cancer cohort.

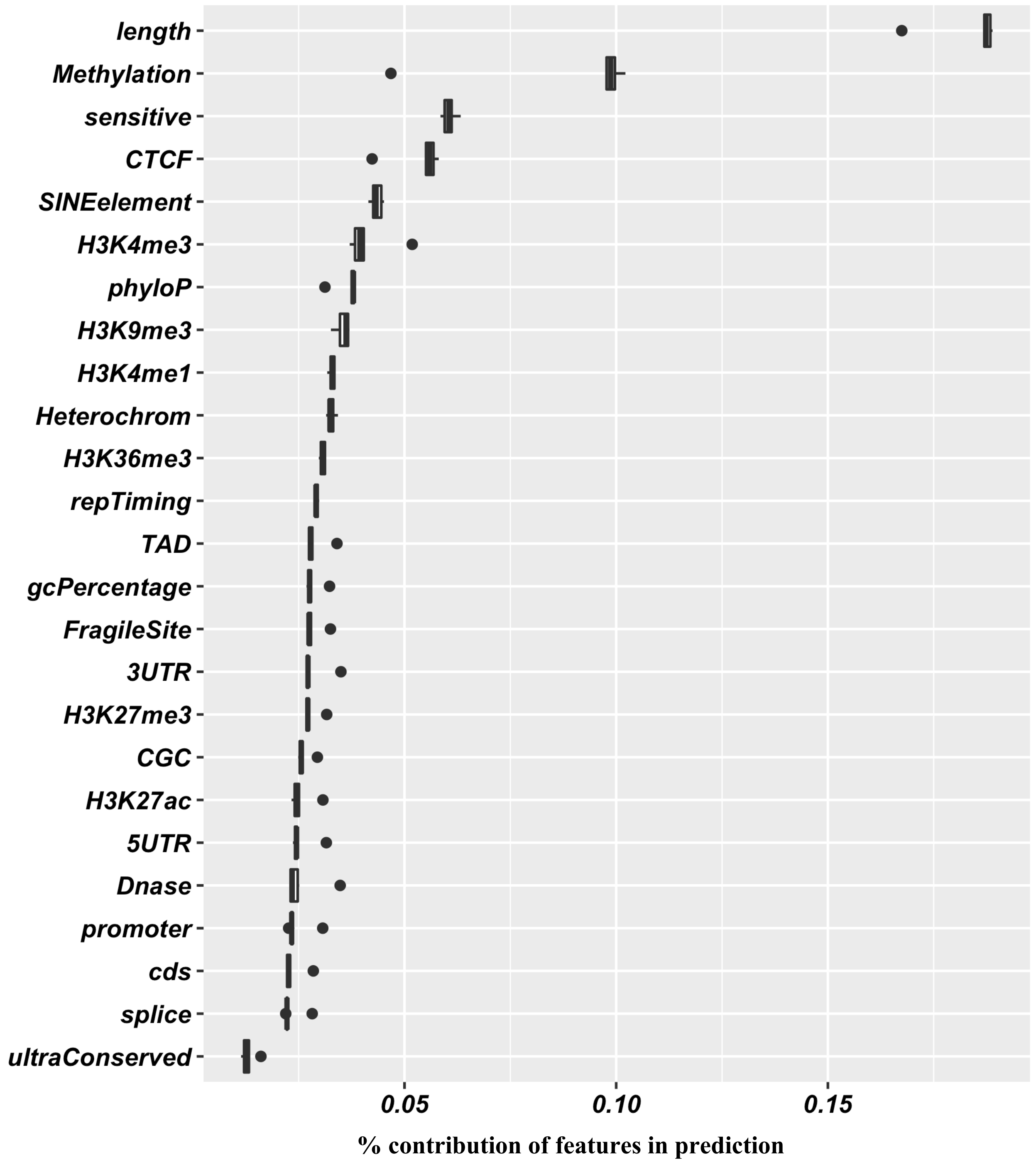


**SI FIG9:** This Figure presents the percentage contribution for distinct features in the overall predictability of germline deletion models for six cancer cohorts. Each data point in the boxplot corresponds percentage contribution of a feature in the overall performance in a cancer cohort.

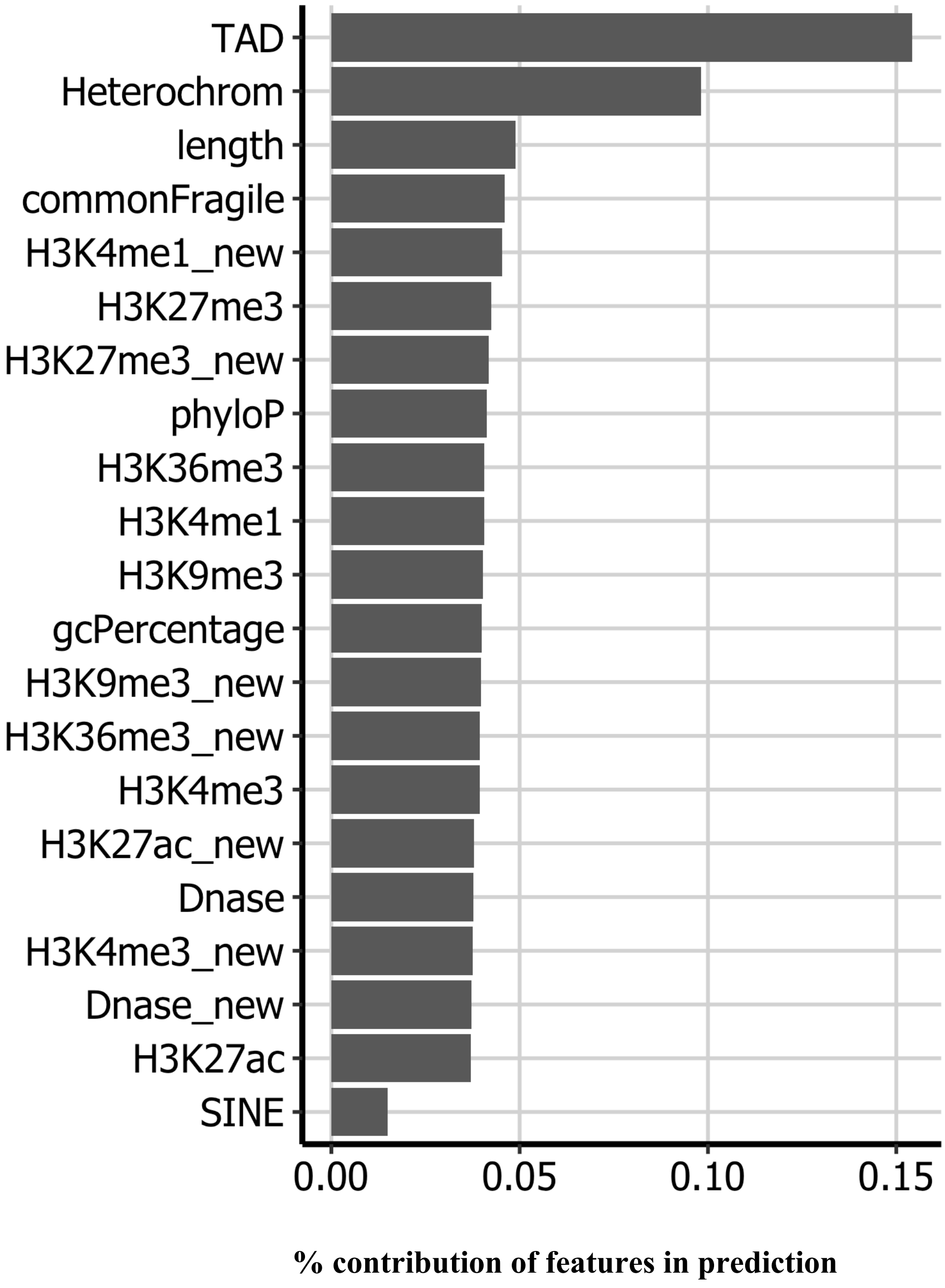


**SI FIG10:** This above bar plot presents the percentage contribution for distinct features in the overall predictability of the germline deletion model for the cardiovascular disease cohort.
